# Characterization of extracellular vesicles produced by *Aspergillus fumigatus* protoplasts

**DOI:** 10.1101/2020.05.21.109926

**Authors:** Juliana Rizzo, Thibault Chaze, Kildare Miranda, Robert W. Roberson, Olivier Gorgette, Leonardo Nimrichter, Mariette Matondo, Jean-Paul Latgé, Anne Beauvais, Marcio L. Rodrigues

**Author notes:** These authors share the senior authorship on this manuscript (Anne Beauvais, Marcio L. Rodrigues). Correspondence to: AB. 25 Rue du docteur Roux, 75015 Paris, France; MLR. Rua Prof. Algacyr Munhoz Mader, 3775, 81350-010 Curitiba/PR, Brazil.

## Abstract

Extracellular vesicles (EVs) are outer membranous compartments produced by yeast and mycelial forms of several fungal species. One of the difficulties to perceive the role of EVs during the fungal life is the fact that an active secretion of these EVs has not been clearly demonstrated in situ due to the presence of a thick cell wall. One alternative to have a better access to these vesicles is to use protoplasts. This approach has been investigated here with *Aspergillus fumigatus*, one of the most common opportunistic fungal pathogens worldwide. Analysis of regenerating protoplasts by scanning electron microscopy and fluorescence microscopy indicated the occurrence of outer membrane projections in association with surface components and the release of particles with properties resembling those of fungal EVs. EVs in culture supernatants were characterized by transmission electron microscopy and nanoparticle tracking analysis. Proteomic and glycome analysis of EVs revealed the presence of a complex array of enzymes related to lipid / sugar metabolism, pathogenic processes, and cell wall biosynthesis. Our data indicate that i) EV production is a common feature of different morphological stages of this major fungal pathogen, and ii) protoplastic EVs are a promising tool to undertake studies of vesicle functions in fungal cells.

**IMPORTANCE:** Fungal cells use extracellular vesicles (EVs) to export biologically active molecules to the outer space. Since fungal cells are encaged in a thick cell wall, it is reasonable to expect that this structure might impact the vesicle-mediated molecular export. In this study, we used protoplasts of *Aspergillus fumigatus*, a major fungal pathogen, as a model to evaluate EV production in the absence of a cell wall. Our results demonstrated that wall-less *A. fumigatus* exports plasma membrane-derived EVs containing a complex combination of proteins and glycans. Our study is the first to characterize fungal EVs in the absence of a cell wall. Our results suggest that protoplasts are a promising model for functional studies of fungal vesicles.

## Introduction

In the *Aspergillus* genus, 90% of all infections resulting in human aspergillosis are caused by *A. fumigatus*, which is the most prevalent mold pathogen in immunocompromised patients (1). *A. fumigatus* has a multifactorial pathogenic arsenal, which allows this organism to successfully establish disease in different hosts (1–4). Aspergillosis begins with inhalation of asexual conidia followed by fungal morphological transition in the absence of a proper immunological response (1).

Fungi, as many other eukaryotic and prokaryotic organisms, produce extracellular vesicles (EVs) (5–8). EVs were first described in the yeast-like pathogen *Cryptococcus neoformans* (9). Subsequent studies demonstrated EV production in yeast forms of *C. gattii*, *Histoplasma capsulatum*, *C. albicans*, *C. parapsilosis*, *Sporothrix schenckii*, *S. brasiliensis, Paracoccidioides brasiliensis*, *P. lutzii, Malassezia sympodialis, Saccharomyces cerevisiae, Pichia fermentans* and *Exophiala dermatitidis* (10–17). In filamentous fungi, the presence of EVs was described in the phytopathogens *Alternaria infectoria* (18) and *Fusarium oxysporum f. sp. vasinfectum* (19), in the dermatophyte *Trichophyton interdigitale* (20), and in the emerging human pathogen *Rhizopus delemar* (21). Recently it was also reported that mycelial forms of *A. fumigatus* produce EVs (22).

A major difficulty to directly address the physiological roles of EVs is the lack of understanding of their intracellular biogenesis and the mechanisms underlying cell wall crossing. An original approach for studying the generation of EVs would be to use protoplasts and look at their active release in the absence of a cell wall. In fact, protoplasts might represent a promising model for the study of EVs, as already suggested by Gibson and Peberdy 50 years ago, who named “subprotoplasts” vesicle-like particles budding from the plasma membrane of *A. nidulans* cells. (23)

Our primary goal in the present work was to search for EVs produced by protoplasts of *A. fumigatus* germinating conidia. Our results revealed the presence of typical EVs in protoplast supernatants incubated under different conditions. EV cargo was directly influenced by the experimental conditions under which *A. fumigatus* was incubated. Our study provides experimental evidence that fungal EVs are produced not only by the mycelial morphological stage of *A. fumigatus*, but also by protoplasts of germinating conidia, especially during cell wall regeneration. To our knowledge, this is the first demonstration that protoplasts may be a useful model to analyze the production and role of EVs in the absence of any cell wall in an experimental setting similar to the ones which allowed the study of exosomes produced by mammalian cells (24).

## Results

### Observation of outer particles resembling EVs in protoplasts of *A. fumigatus* germinating conidia

To experimentally overcome the difficulties of detecting EVs due to the presence of a thick cell wall, we adopted an experimental model using protoplasts. Fungal cells lacking cell walls, which were obtained by enzymatic digestion, can reconstruct their walls in osmotically stabilized media (25–29). In this context, we first compared the morphological aspects of *A. fumigatus* freshly obtained protoplasts with cell-wall regenerating cells by scanning electron microscopy (SEM). Under both experimental conditions, we observed ~50 nm round-shaped extracellular structures in an apparent association with the fungal surface (**Figure 1**). When the protoplasts were incubated under conditions of cell wall synthesis, the outer particles were more numerous. During regeneration, polysaccharide neosynthesized were seen above the vesicles (data not shown).

**Figure 1.**
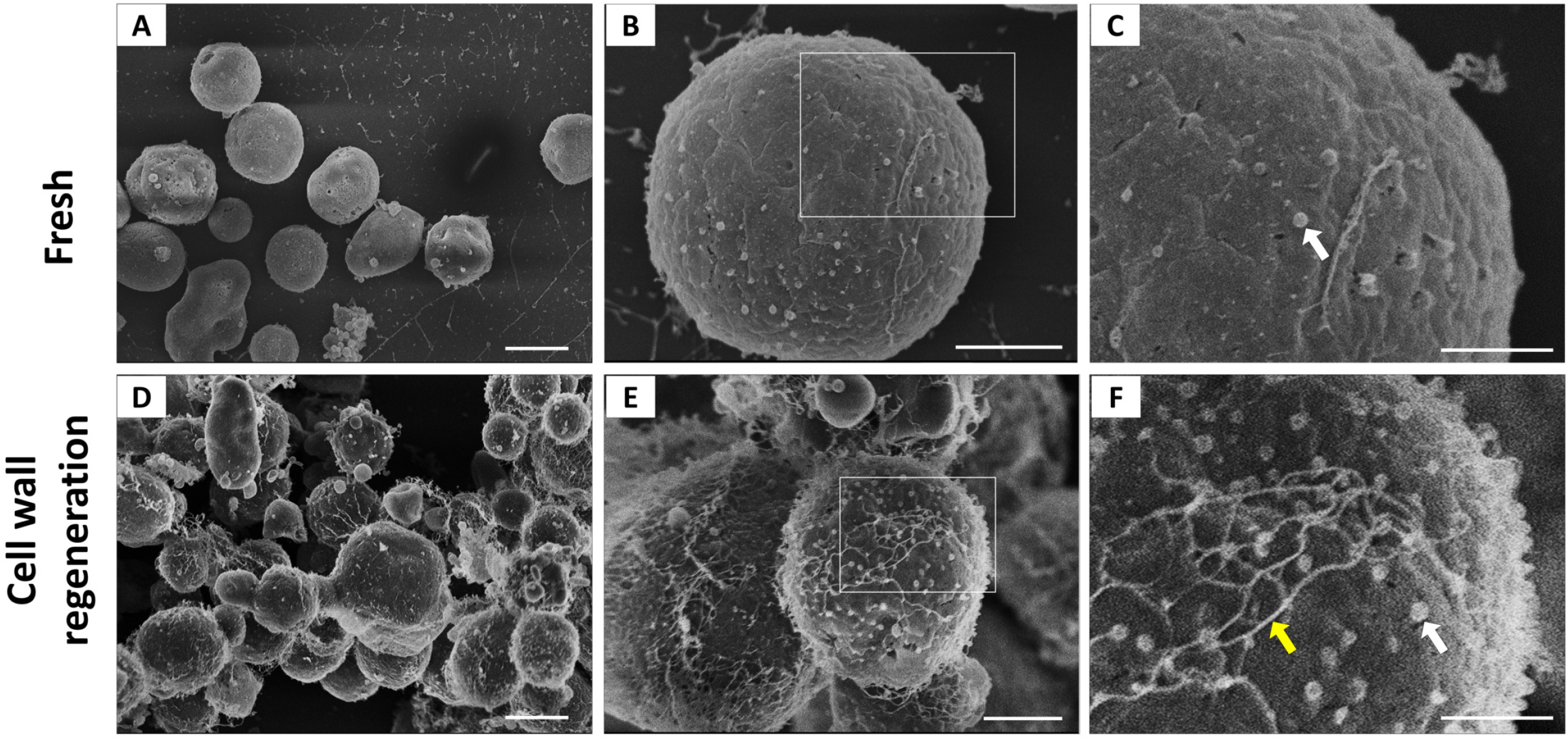
Morphological aspects of freshly prepared and cell wall regenerating protoplasts. Fresh protoplasts (A-C) and cell wall regenerating cells (D-F) are shown under conditions of increasing magnification by SEM. C and F represent magnified views of the boxed areas in B and E, respectively. Magnified views suggested the occurrence of outer particles with properties compatible with EVs (white arrows). Under cell wall regenerating conditions, a fibril-like network was more abundantly detected (yellow arrow). Scale bars represent 5 μm in panels A and D, 2 μm in panels B and E and 1μm in panels C and F.

To better analyze these vesicle-like particles, we checked the membrane distribution of freshly prepared protoplasts and protoplasts incubated under conditions of stimulation of cell wall synthesis or no synthesis of the cell wall. Incubation outcomes were monitored by staining the protoplasts with an anti-glucan antibody and observation by fluorescence microscopy. Membranes were stained with the lipophilic dye DiI. *A. fumigatus* protoplasts manifested the complex membrane distribution that is typically observed in most eukaryotic cells (**Figure 2A**). As expected, glucan was detected at the background levels in freshly prepared protoplasts and protoplasts incubated in KCl alone. Under conditions of cell wall synthesis, surface glucan was unequivocally detected. A detailed analysis of the relationship between membrane staining and glucan detection in these cells revealed several regions of membrane projection to the outer space (**Figure 2B**). The projected regions were closely associated with glucan detection, and, in fact, the polysaccharide was apparently surrounded by the membranous compartments (**Figure 2C**). The occurrence of membrane projections in protoplasts incubated under conditions of cell wall synthesis was confirmed by super-resolution SEM (**Figure 2D**).

**Figure 2.**
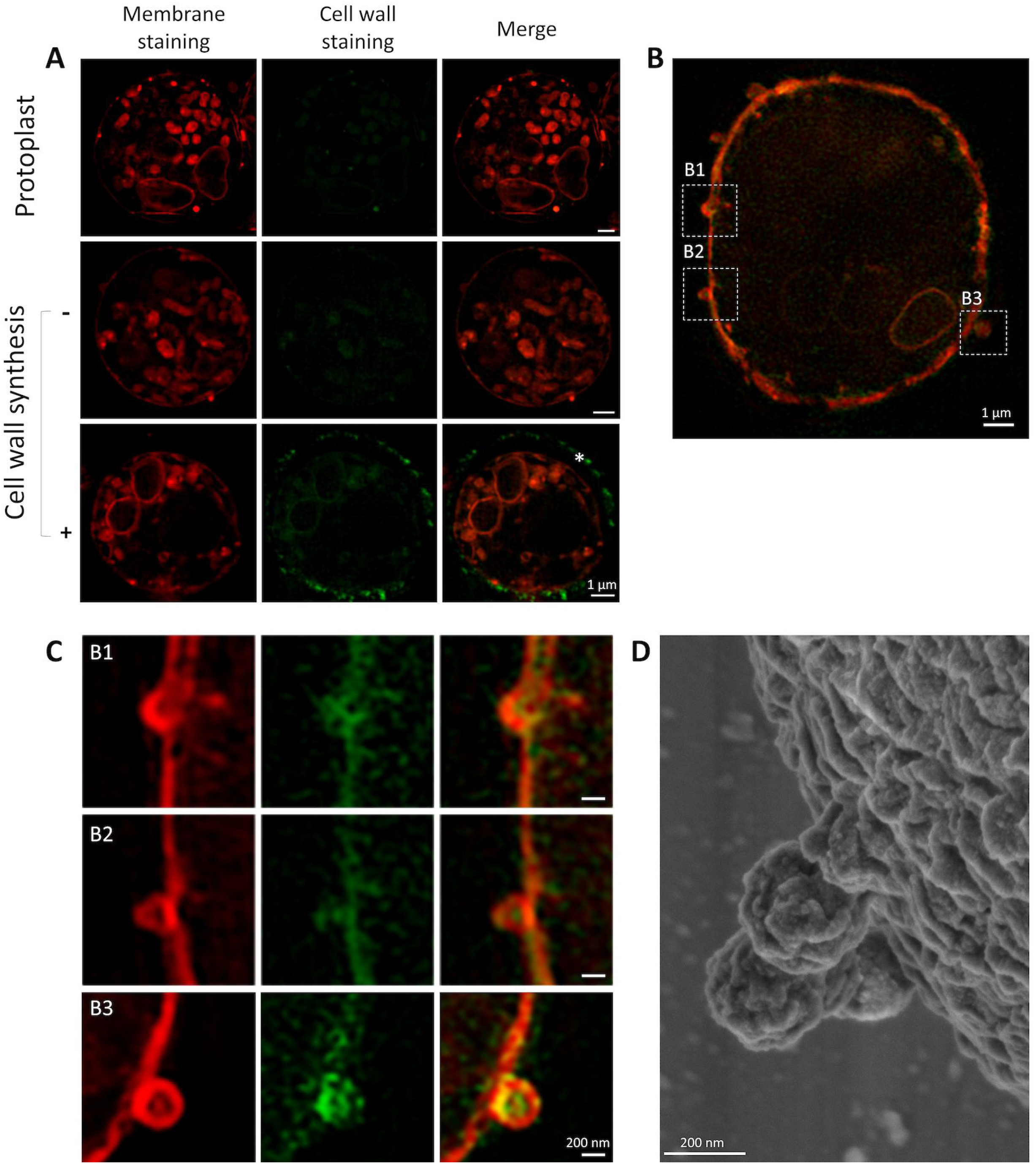
Membrane projections in *A. fumigatus* protoplasts. A. Freshly purified protoplasts were stained with DiI, a lipophilic die (red fluorescence). Cell wall staining with an anti-glucan antibody was at the background levels. Similar results were observed for protoplasts incubated under non-regenerating conditions. During cell wall regeneration (2 h), glucan staining (green fluorescence) was abundant at the cell surface. A detailed analysis of these cells (B) revealed an association between glucan staining and outer membrane projections in cell wall regenerating protoplasts after 90 min of incubation, as evidenced in the boxed areas (B1, B2 and B3), which were amplified and shown in C. A detailed view of the surface of protoplasts by super-resolution SEM (D) confirmed the occurrence of outer particles budding out of the plasma membrane.

### Protoplasts of *A. fumigatus* germinating conidia release EVs

To analyze the occurrence of EVs, protoplasts were adjusted to 10^8^ protoplasts/ml, and cell viability was in the 87-93% range during all the duration of the experiments. Supernatants obtained from protoplasts incubated under regenerating and non-regenerating protoplasts were fractionated by ultracentrifugation, and the resulting pellets were analyzed by transmission electron microscopy (TEM). Membranous structures with a typical bilayer, morphological aspects, and dimensions previously observed for fungal EVs were isolated from protoplast supernatants (**Figure 3A, B, E and F**). Regenerating protoplasts produced EVs that were apparently associated with fibrillar material (**Figure 3G and H**). Most of the vesicles were in the 200 nm diameter range, and this visual perception was confirmed by NTA, which revealed a peak of vesicle detection at this size range (**Figures 3I and J**). EVs obtained from control or regenerating protoplasts had very similar diameter distribution profiles.

**Figure 3.**
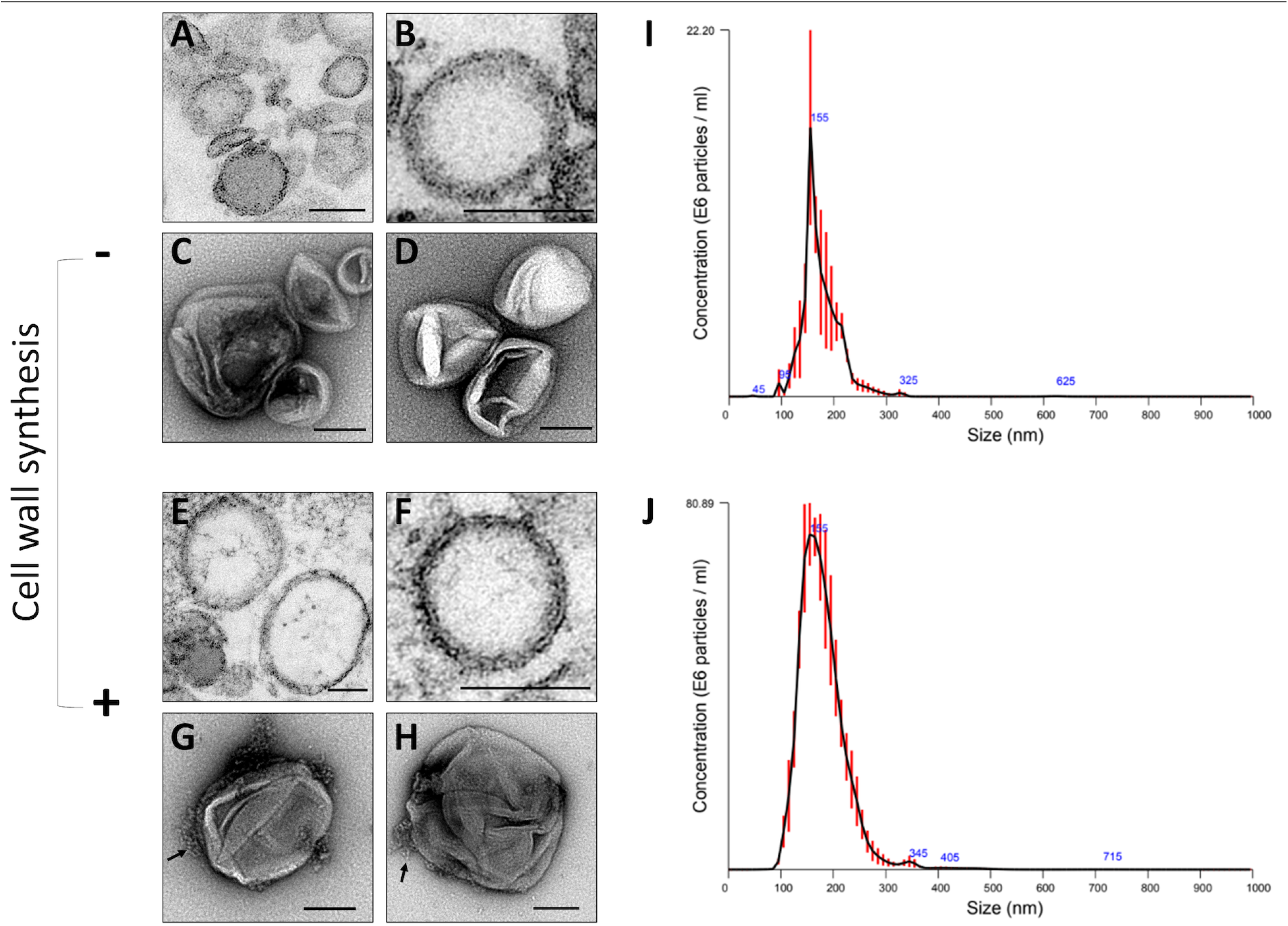
Analysis of EVs obtained from *A. fumigatus* protoplasts. Protoplast EVs were analyzed by regular TEM (A, B, E, F) or after negative staining (C, D, G, H). Independent illustrations of each condition are shown for each technique. Under conditions stimulating cell wall synthesis, fibril-like structures associated with EVs were observed (G-H, arrows). Scale bars correspond to 100 nm. NTA of isolated vesicles (I, J) demonstrated a similar distribution of EVs in the 50-300 nm diameter range, independently on the condition of incubation of the protoplasts.

### EVs are more abundantly detected in the supernatants of regenerating protoplasts

We quantified EV production in the experimental systems explored in our study by different approaches. First, independent replicates were submitted to quantitative NTA. This analysis revealed that the EV/cell ratios were at the background levels when the samples from freshly prepared protoplasts were analyzed (**Figure 4A**). Even though some EVs were released over time in the supernatants of control protoplasts, their number was drastically increased in the supernatant of regenerating protoplasts. The NTA data agreed with the quantification of the sterols in the EV-containing supernatants (**Figure 4B**). This increase in the sterols was not associated with an enhancement in the total amount of the protoplast sterols occurring during cell wall regeneration (data not shown).

**Figure 4.**
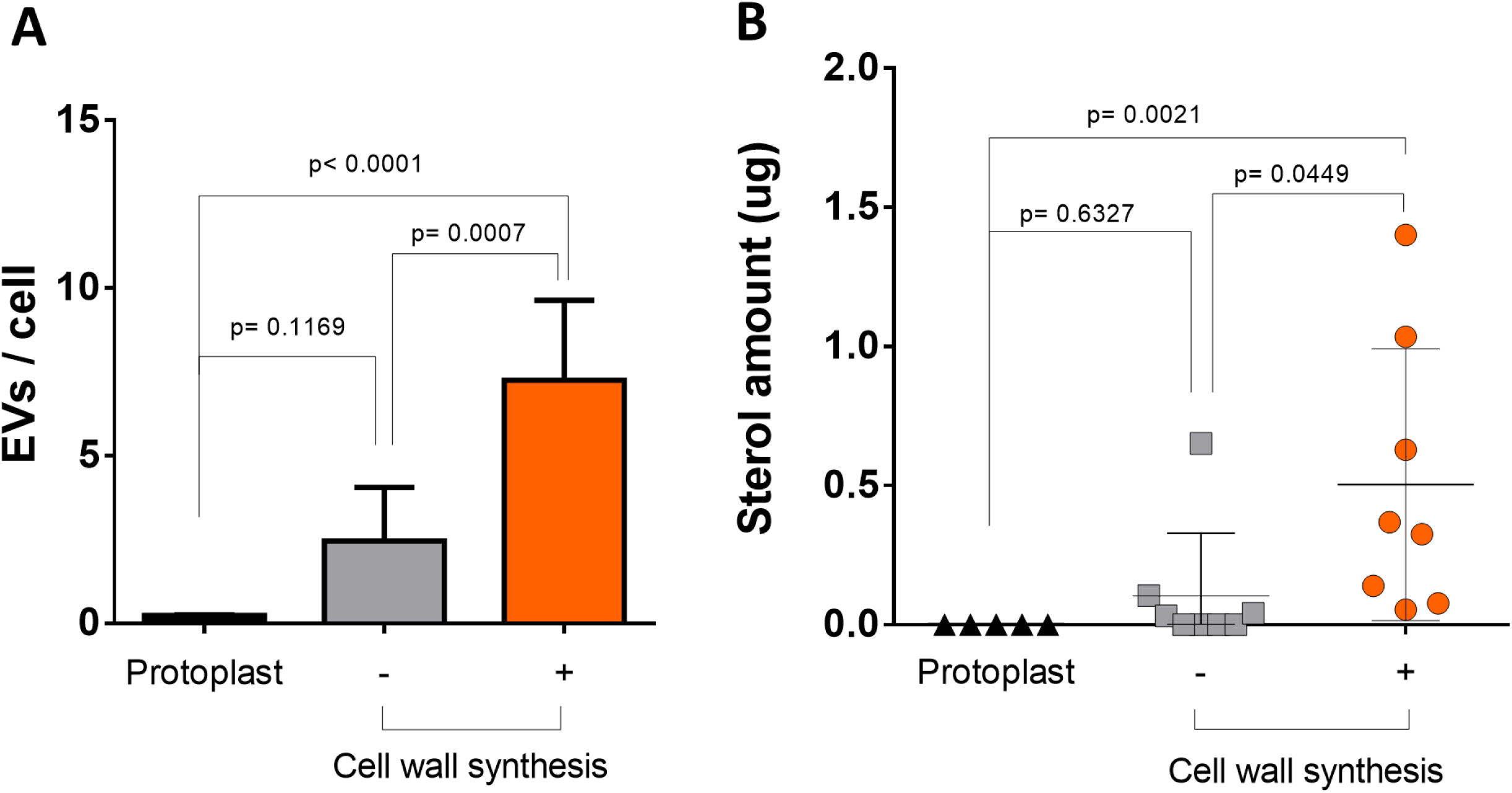
EV quantification during the cell wall synthesis process in *A. fumigatus* protoplasts. A. Quantitative NTA of EVs produced by freshly prepared protoplasts and from protoplasts incubated under conditions of cell wall regeneration or non-regeneration. B. Determination of sterol concentration in EVs obtained from supernatants of fresh protoplasts, non-regenerating protoplasts, and protoplasts incubated under cell wall regenerating conditions. In A and B, values are reported as means ± standard deviations obtained from at least two and five independent experiments, respectively.

### Glycan components of *A. fumigatus* EVs

The molecular composition of the EVs released by the protoplasts was investigated. Since GAG is a maker of polysaccharide secretion by the *A. fumigatus* mycelium (4), we first showed the presence of this polysaccharide during protoplast regeneration (**Figure 5A**). We then proved that GAG was present in EVs isolated from protoplasts incubated under the conditions of cell wall regeneration and absent in control vesicles (**Figure 5B**). These results agreed with the compositional analysis of the carbohydrate units of EVs. Glucosyl (Glc), mannosyl (Man), and galactosyl (Gal) units were detected in EV preparations obtained from both cell wall regenerating and non-regenerating protoplasts (**Figure 5C**). N-acetyl-galactosaminyl (GalNAc) residues, which are markers of GAG, were only found in regenerating protoplasts. In addition, Glc was significantly increased in EV samples from these cells, compared to non-regenerating protoplasts (p = 0.004). N-acetyl-glucosamine (GlcNAc) residues were absent in all samples.

**Figure 5.**
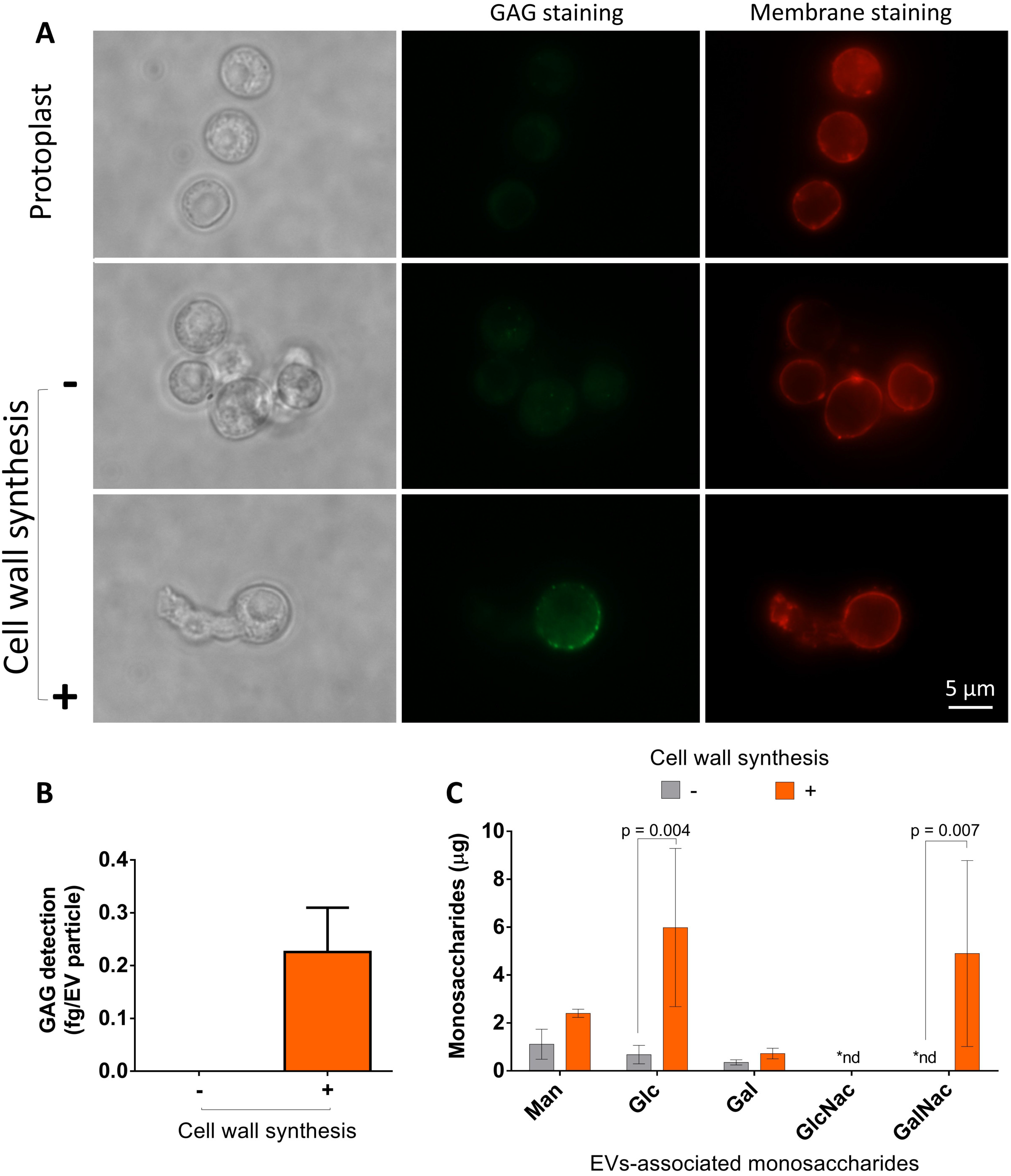
Analysis of glycan synthesis during cell wall regeneration in *A. fumigatus* conidial protoplasts. **A**. Membrane and GAG staining in *A. fumigatus* protoplasts. All cells were efficiently stained with DiI (red fluorescence). During cell wall synthesis, GAG was detected in association with the fungal surface. Scale bar corresponds to 5μm B. Serological detection of GAG (ELISA) in EVs obtained from protoplasts. Positive reactions with a GAG-binding antibody were only observed in EVs obtained during cell wall synthesis. C. GC-MS analysis of sugar units of EVs. In agreement with an involvement of EVs with cell wall synthesis, GalNAC (a GAG component) was only observed in EVs obtained from protoplasts during cell wall regeneration. The increased detection of Glc during cell wall synthesis is consistent with EV-associated glucans.

### Proteomic analysis of EVs

Proteomic analysis revealed only 142 proteins in EVs produced by fresh protoplasts, contrasting with the detection 2056 proteins in vesicles from regenerating protoplasts (Supplemental Table 1). All the 142 EV proteins detected in fresh protoplasts were found in samples obtained from cell wall regeneration conditions. Although a lot more EVs were produced in the regenerating protoplasts, the qualitative protein composition of the non-regenerating protoplast was similar (Supplemental Table 1).

The predicted GO classification of all EV-related proteins identified numerous terms (**Figure 6**). As previously described for several fungal EVs (11, 14, 22, 30), the shared GO terms corresponded to proteins involved in a wide range of processes of fungal physiology. Most of the biological processes (680 GO terms) were shared by the regenerating and non-regenerating conditions, but 50 GO terms of them were specifically found in the cell wall regenerating system. Using the Uniprot (https://www.uniprot.org/) and AspGd (http://www.aspgd.org/) databases, we classified the EV composition according to the *A. fumigatus* cell wall assembly process. The cell-wall associated proteins identified in EVs obtained from each condition are listed in Table 1. We detected several proteins related to (i) cell wall synthases, including the β1-3 glucan synthase Fks1 and its GTPase activator Rho1 (31), α1,3 glucan synthases Ags (32), chitin synthases (33), Ktr mannosyltransferases involved in the synthesis of galactomannan (34) and enzymes belonging to the GAG biosynthetic pathway (Ugm1, Gt4c) (35); (ii) cell wall remodeling enzymes, including Gel and Bgt glucosyltransferases involved in the elongation and branching of the β1-3 glucan (36); (iii) some enzymes involved in mannosylation, including the multi enzymatic complexes mannan polymerase I and II involved in the synthesis of one of the two conidial mannan (Mnn proteins, Van1, Och1) (37), and Pmt O-mannosynltransferases involved in the 0-mannosylation of cell wall remodeling enzymes (38). Other cell wall related proteins, including Mp1, PhiA, MidA, GPI-anchored proteins including Ecm33, mannosyltransferases (Alg2, Och1and Mnn2), and the putative glycan biosynthesis protein Pig1 were also detected.

**Figure 6:**
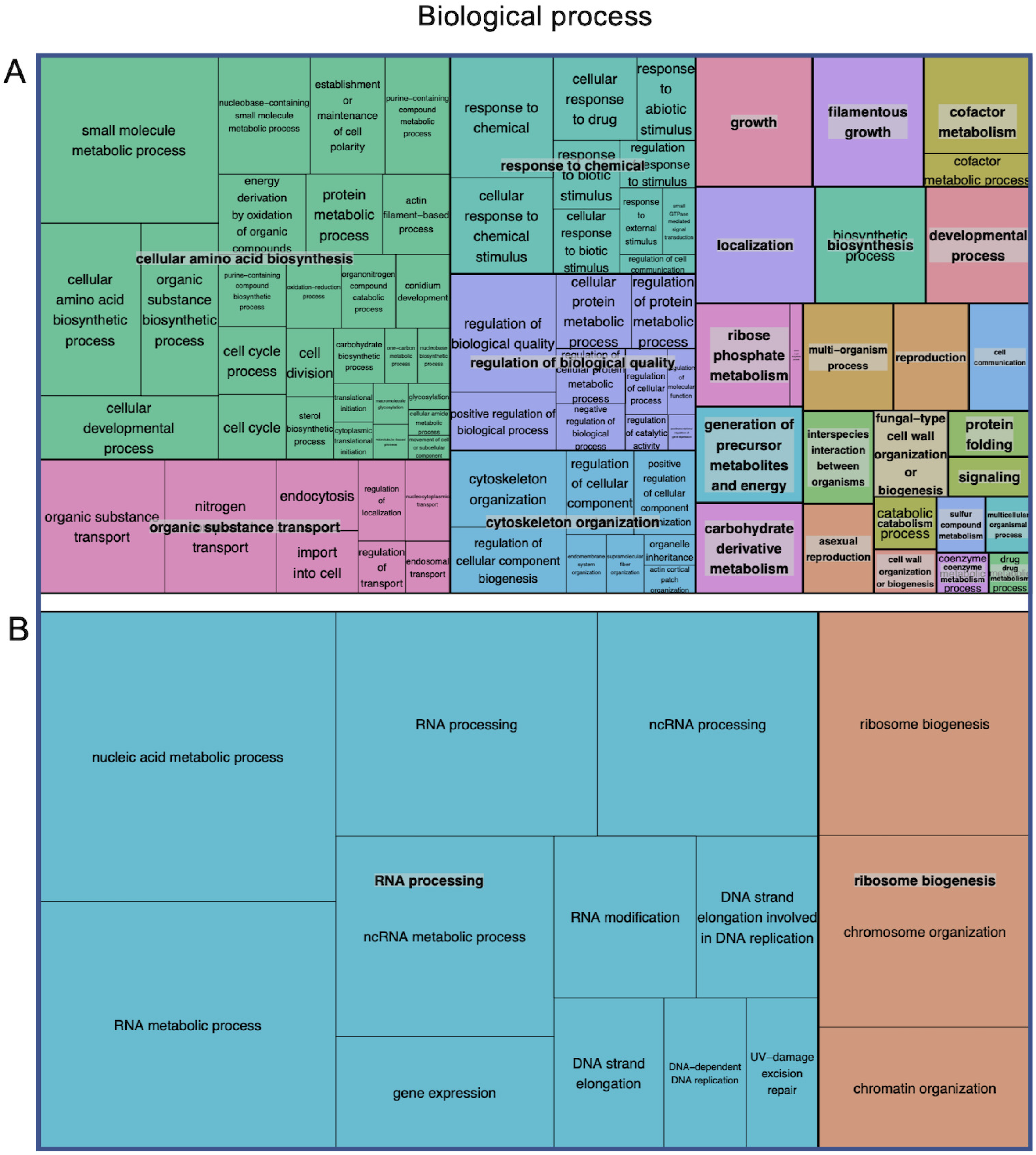
Proteomic analysis of EVs obtained from supernatants of *A. fumigatus* protoplasts. TreeMap view of all biological processes, cellular components and molecular functions with which vesicular proteins were associated. Panel A shows the biological processes common to regenerating and non-regenerating conditions. Panel B shows the processes that were exclusively found under conditions of cell wall regeneration. Rectangle areas reflect the p-value of enrichment of GO terms in the *Aspergillus* database. GO terms are gathered under summarized terms using the REVIGO tool (71).

**Table 1:**
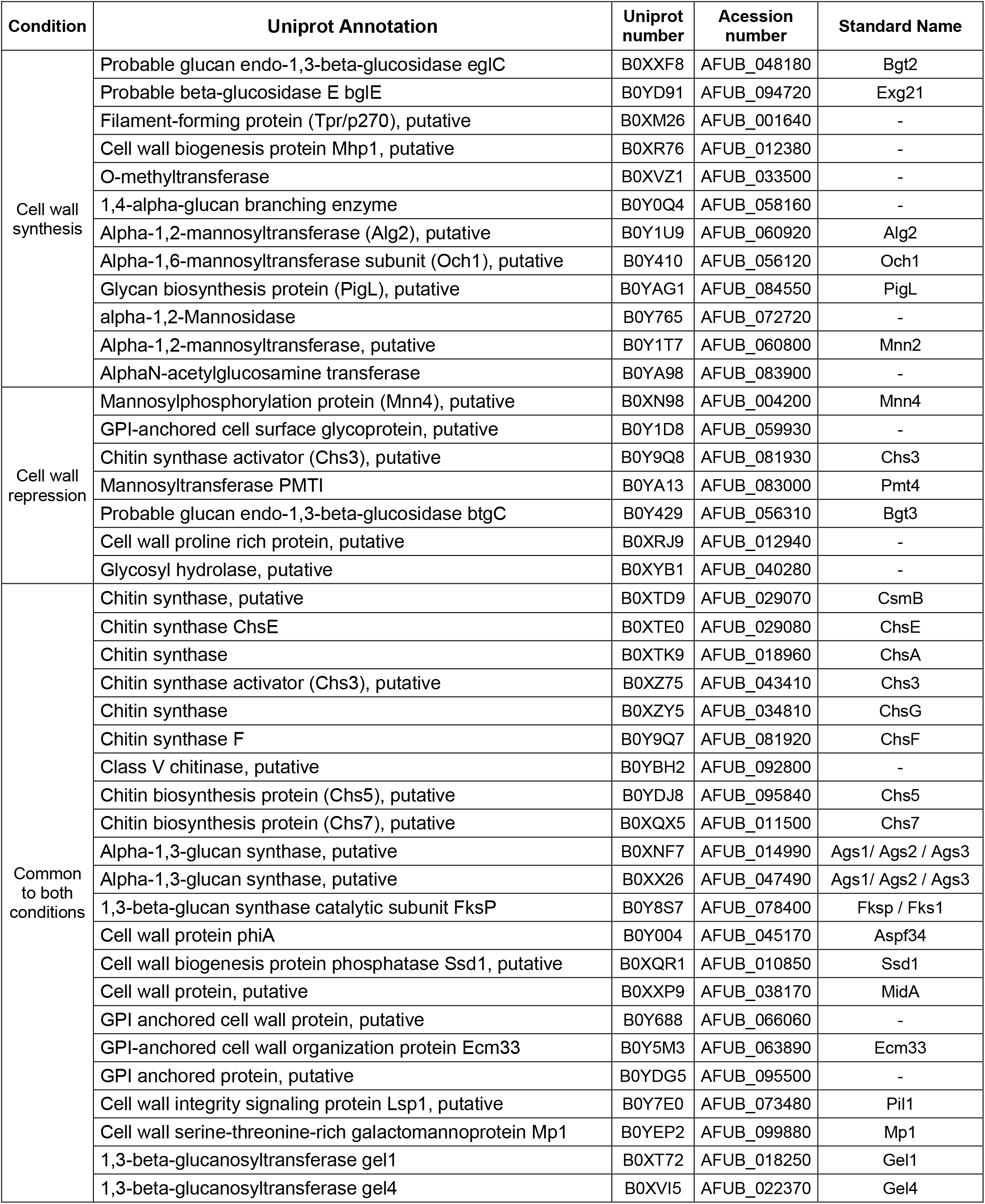
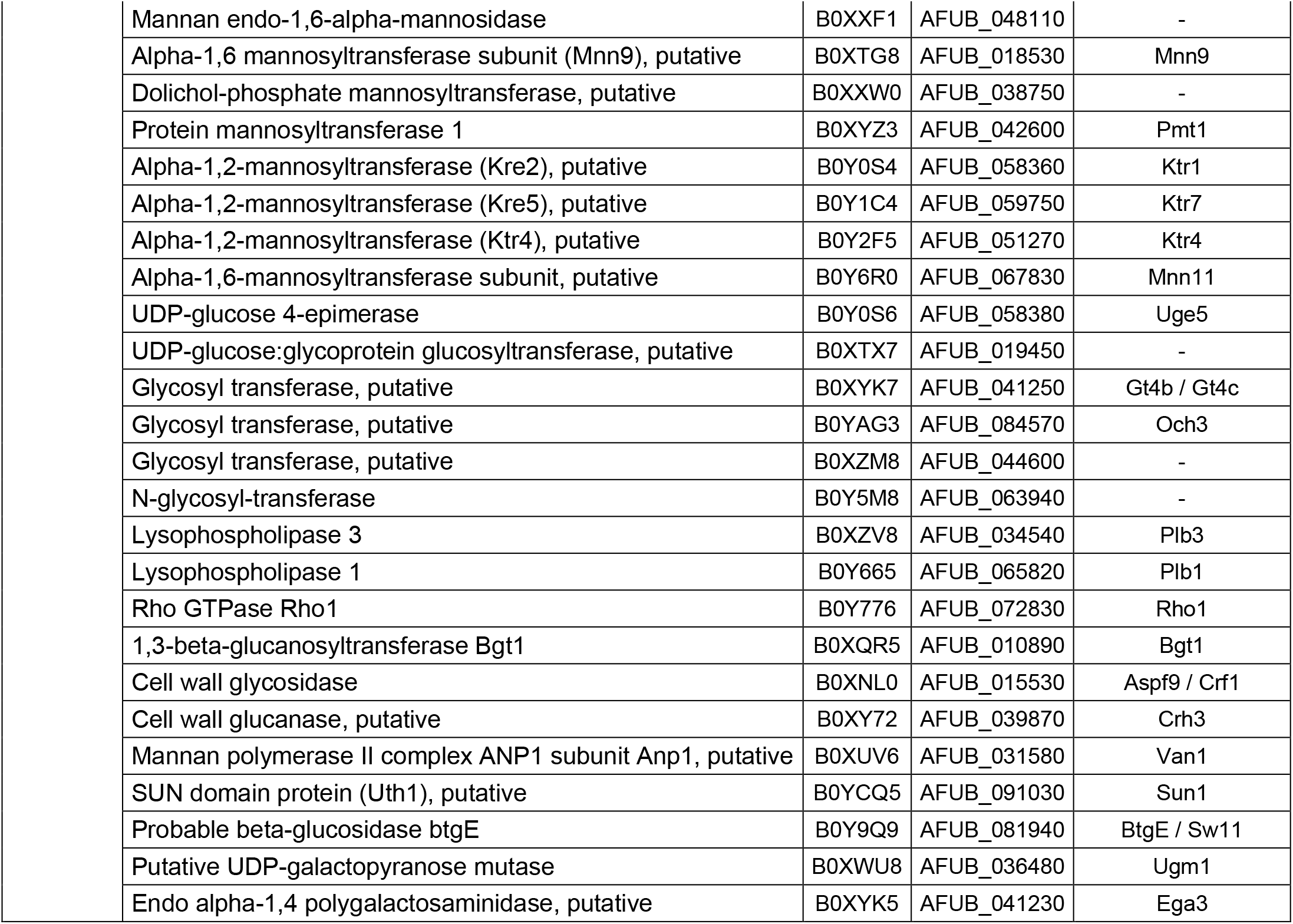
Cell wall-associated proteins found in *A. fumigatus* EVs produced by protoplasts.

## Discussion

Yeast forms of different fungal species produce outer membrane structures classified as EVs (39). More recently, it has been demonstrated that filamentous forms of fungi also produce EVs (18–20, 22). Although the functional impact of these findings is still not clear, they confirm that EVs are released by different morphological stages of fungi as part of distinct physiological events. One of the difficulties of understanding the physiological function of EVs is the presence of a cell wall surrounding the cell protoplasm. Our study has shown that the use of alive wall-less stages like protoplasts can alleviate this problem and that these cells are suitable for the study of EV release. Of note, protoplasts of other species, including *C. albicans, S. pombe*, and *N. crassa*, also produced extracellular particles resembling EVs (23, 26, 40), as concluded from early microscopic analysis of fungal cells.

Our analysis of protoplast forms of *A. fumigatus* germinating conidia by SEM revealed the presence of particles with general properties compatible with those of EVs being released from the plasma membrane. These vesicular particles were morphologically similar to those observed in association with the cell wall of *C. neoformans* (41). This result and the observation of vesicles emerging from the *A. nidulans* surface (23) suggested that *A. fumigatus* protoplasts are efficient producers of EVs.

EVs released by regenerating protoplasts showed a fibril-like material attached to the lipid surface, as evidenced by TEM of isolated EVs. Even though EVs were produced in the highest number in regenerating protoplasts, it is noteworthy that EV release was not uniquely associated with cell wall biosynthesis since the non-regenerating protoplasts also produced EVs. This observation suggests that EV release is not exclusively related to the synthesis of the cell wall and that the release of EVs in non-growing cells may be also a response of the fungus to extracellular stress such high osmotic pressure or lack of nutrients. Noteworthy, EVs from cell wall regenerating cells and non-regenerating protoplasts differed in relative concentration. This observation could simply correspond to a more efficient metabolic response in cell wall-synthesizing cells, but also to a possible association of EVs with cell wall synthesis in *A. fumigatus* protoplasts. This last supposition remains to be experimentally proved.

Our carbohydrate analysis revealed the presence of Man and Gal, and an increased amount of Glc and the presence of GalNAc in EVs obtained from regenerating protoplasts. Glc is a marker of β or α1,3 glucan. Man and GaI are markers of galactomannan. β or α1,3 glucan are likely incorporated in the EVs during their formation, as concluded from the previous demonstration that cell wall polysaccharides are synthesized on the internal side of the plasma membrane level and extruded in the cell wall at the C terminal pore-like part of the respective enzymes (42, 43), as also shown in Figure 2 of our study. Galactomannan is assembled in the Golgi apparatus and secreted to the plasma membrane before being cross-linked to β1,3 glucan supposedly by extracellular transglycosidases (35). Therefore, the presence possible of glucans and galactomannan in EVs could be a consequence of their original association with the plasma membrane. GAG, which is a virulence-associated component of the *A. fumigatus* extracellular matrix (44, 45), is localized on the surface of the cell wall, where it acts as a component of the fungal extracellular matrix (46). The detection of GalNAc residues only in EVs obtained from regenerating protoplasts suggests that GAG is transported by vesicles through the cell wall to be deposited on the cell surface. Alternatively, GAG could be loosely associated with EVs, considering its “sticky” nature due to its great ability to form unspecific hydrogen bonds. Similar observations were described in *C. neoformans*, in which EVs contained the extracellular polysaccharide glucuronoxylomannan (47).

It is still unknown whether the EVs characterized in our protoplast model represent the vesicular structures produced by intact *A. fumigatus*. Although the general properties of protoplast (this study) and mycelial EVs (22) are similar, we identified a higher diversity of EV proteins in comparison to *A. fumigatus* mycelial vesicles (22). It is important to highlight, however, that the strains used in these studies were distinct. In addition, all studies on fungal EVs produced so far used regular cultures for vesicle isolation and not regenerating protoplasts. Compositional differences between the current study and previous reports are, therefore, expected. Thirty two out of the 60 proteins described for the mycelial *A. fumigatus* EVs were also found in our study. Since protoplasts are produced from the germ tube tips, they contained all the cell wall synthesis and remodeling machinery normally present in the plasma membranes of fungal apexes (35, 48, 49). Consequently, it is not surprising that protoplast EVs contained many of these cell wall enzymes. Some of these proteins, including Fks1, CsmB and Chs,Gel4, Pmt4, Ktr4, Ktr7 and Gt4C, are essential for the synthesis of the major cell wall polysaccharides β and α1,3 glucans, chitin, galactomannan and GAG, and for branching/elongation of the β1,3 glucan in *A. fumigatus* (50–52). The absence of GlcNAc (N-acetyl-glucosamine residues) may suggest that chitin-related molecules are absent in the EVs. However, we cannot rule out the possibility that a slower kinetics of chitin synthesis affected our experimental model. For instance, preliminary experiments of polysaccharide immunolabeling of regenerating protoplasts showed that β1,3 glucan was the first polysaccharide detected on the surface of the protoplasts, following by α1,3 glucan. Chitin was the last polysaccharide to be detected (A. Beauvais, unpublished results).

Proteins that are not predicted to be in the extracellular milieu were abundantly detected in EVs from *A. fumigatus* protoplasts. This observation agrees with numerous reports on the protein composition of fungal EVs (8, 11, 14, 30) and with the fact that these membranous structures have their biogenesis linked to the cytoplasm and the plasma membrane (53). Additionally, it was suggested that the cell wall is a storage site for many fungal proteins (54–57). Cell wall-related proteins, including glucanases, phiA, Ecm33, Gel1, and Gel4, were also identified in *A. fumigatus* culture fluids (57).

Our current results contribute to a better understanding of the properties of fungal EVs. To our knowledge, this is the first characterization of EVs in protoplasts obtained from germinating conidia. Our results suggest that these cellular forms represent a promising model to explore novel roles of fungal EVs in many fungal species. In the *A. fumigatus* model, phagocytic cells stimulated with EVs increased their ability to produce inflammatory mediators and to promote fungal clearance (22). These results support the relevance of using protoplastic fungal EVs devoid of any cell wall contaminants to better understand their role in both physiology and immunopathogenesis of *A. fumigatus*.

## Acknowledgments

M.L.R. was supported by grants from the Brazilian Ministry of Health (grant number 440015/2018-9), Conselho Nacional de Desenvolvimento Científico e Tecnológico (CNPq, grants 405520/2018-2, and 301304/2017-3) and Fiocruz (grants VPPCB-007-FIO-18 and VPPIS-001-FIO18). The authors also acknowledge support from the Instituto Nacional de Ciência e Tecnologia de Inovação em Doenças de Populações Negligenciadas (INCT-IDPN). J.R. developed part of this work as a Ph.D student supported by FAPERJ (E-26/201.991/2015) and is currently a postdoctoral fellow at Institut Pasteur (Paris, France), funded by the CAPES-Cofecub Program (88887.142840/2017-00). We are grateful to Leandro Honorato (Instituto de Microbiologia Paulo de Goes, UFRJ, Brazil) and Luna Sobrino Joffe (Stony Brook University, USA) for help with protoplast purification, Flavia Reis (Fiocruz, Brazil) for help with NTA, Fernando P. de Almeida for the help with super resolution microscopy (CENABIO/ UFRJ, Brazil), and Vishukumar Aimanianda (Institut Pasteur, France) for discussions and for providing some of the antibodies for this study. This research was partly funded to JPL by the Laboratoire d’Excellence “Integrative Biology of Emerging Infectious Diseases” (grant n°ANR-10-LABX-62-IBEID), la Fondation pour la Recherche Médicale (DEQ20150331722 LATGE Equipe FRM 2015).

## Material and Methods

### Growth conditions and preparation of *A. fumigatus* protoplasts

The *A. fumigatus* reference strain used in this study was CEA17ΔakuB^KU80^ (ku80), which is deficient in non-homologous end joining (58). The CEA17ΔakuB^KU80^ strain was conserved on 2% (w/v) malt agar slants. Five days-old conidia were recovered from the slants by vortexing with a 0.05% (v/v) aqueous Tween 20 solution and filtered through a 40 μm cell strainer. For protoplast preparation, 7 x 10^9^ conidia suspended in 0.05% Tween were centrifuged at 3,000 x *g* for 10 minutes. The supernatant was discarded, and the cells were suspended in 10 ml of sterile water. The conidia were inoculated in 600 ml of germination medium (1% yeast extract, 3% glucose, and 0.6M mannitol) and then incubated with shaking for 14 h at 30°C. Germinated conidia were harvested and separated from dormant conidia by filtering the sample through a sterile miracloth-lined funnel and then washed with 200 ml of sterile osmotic medium (OM; 1.2 M MgSO4.7H_2_O, 0.09 M K_2_HPO_4_, and 0.01M KH_2_PO_4_, pH 5.8), two-fold diluted. After washing, the germinated conidia were suspended in 20 ml OM containing Glucanex^®^ at 30 mg/ml (Novo Nordisk Ferment Ltd., catalog number CH4243). The cells were gently homogenized, and the suspension was adjusted to a final volume of 250 ml with a filtered-sterile Glucanex^®^ solution for hydrolysis of the cell wall.

Cell wall digestion was performed in 2 l Erlenmeyer flasks for 2 h at 37°C, with gentle shaking (60 rpm) until sufficient protoplasts were released, as assessed microscopically. For protoplast recovery, the cells were harvested by filtration in a sterile glass Buchner funnel (porosity 2). After filtration, two volumes of sterile 0.3 M KCl were added to 1 volume of the protoplast suspension in Glucanex^®^. The mixture was centrifuged at 5,000 x *g* (20 min, 25°C). The pellet was suspended in sterile 0.6 M KCl and washed twice (3,000 x *g*, min/min, 10 min, 25°C) to eliminate the remaining Glucanex^®^. The final pellet of fresh protoplasts was divided into two equal parts. Each part was suspended in 10 ml of sterile 0.6 M KCl and further incubated under nonregenerating or cell wall synthesis conditions. The viability of the protoplasts was assessed by using trypan blue dye exclusion at different time points, from 0 to 2 h.

For cell wall regeneration, the protoplasts were incubated in 400 ml of a minimal medium (MM) supplemented with 0.6 M KCl for 2 h, at 37°C under shaking (120 rpm). The MM was prepared as previously described with some modifications (59, 60) and contained 1% (wt/vol) glucose, 20 mM glutamine, 0.052% KCl, 0.052% MgSO_4_.7H_2_O, 0.152% KH_2_PO_4_ and 1 ml trace-element solution, pH 6.5. Alternatively, protoplasts were incubated for 2 h in a non-regenerating solution of 0.6 M KCl or immediately processed for the analyses described below.

### Microscopic analysis of protoplasts

Freshly purified protoplasts, as well as protoplasts obtained from cell wall regenerating or non-regenerating conditions, were fixed with 2% formaldehyde in 0.6 M KCl and stored at 4°C. For fluorescence or super-resolution microscopy, the cells were washed twice with PBS and blocked with the superblock blocking buffer in PBS (Thermo Scientific, catalog number 37515) for 1 h at 37°C. Surface α-(1,3)-glucan of protoplasts was labeled with the MOPC 104E monoclonal antibody (2 μg/ml, 1 h at 37°C) (61). Alternatively, the cells were stained with an anti-GAG monoclonal antibody (20 μg/ml, 1h at 37°C) (44). After two washes in PBS, each preparation was incubated with the appropriate secondary antibodies (anti-mouse IgM Alexa Fluor 488 for glucan staining; anti-mouse IgG Alexa Fluor 488 for GAG; both diluted 1:100 in blocking buffer). After incubation for 1 h at 37°C, the cells were washed three times with PBS. The protoplast membranes were finally stained with the Vybrant™ Dil Cell-Labeling Solution (Molecular Probes, catalog number V22885) at 5 μM (30 min, 37°C) and washed one final time with PBS. The cells were placed on glass slides covered with the ProLong^®^ Gold antifade reagent. The cells were microscopically observed under the Zeiss Axioplan 2 fluorescence microscope or the Zeiss Elyra PS.1 super resolution microscope under structural illumination mode. Images were obtained with the ZEN 2.1 Software.

Super resolution scanning electron microscopy (SEM) was performed as previously described (62). Briefly, 10^6^ protoplasts were fixed on sterile glass coverslips (previously coated with poly-L-lysine) overnight at 4°C on 24 well plates. The samples were dehydrated in graded ethanol series (30%, 50% and 70% for 5 min and 95% and 100% for 10 min), critical point dried in CO_2_, mounted on stubs and coated with carbon. Observation of the protoplast cell surface was performed with an Auriga 40 field-emission scanning electron microscopy (FE-SEM) microscope (Zeiss, Germany).

Protoplasts were also were analyzed with a JEOL JSM-6700F apparatus, which is an ultra-high-resolution field emission scanning electron microscope equipped with a cold-field-emission gun and a strongly excited conical lens. The secondary-electron image resolution was 1 nm at 15 kV and 2.2 nm at 1 kV. Pieces of culture were frozen using a Gatan Alto 2500 cryo-stage and cryo-preparation chamber. The preparation conditions were as described in Paris et al. (63).

### EV isolation and physical-chemical analysis

The isolation of EVs from protoplast supernatants was performed as previously described for yeast cells (39), with minor modifications. Briefly, after each incubation period, the supernatants were separated from the protoplast cells by centrifugation at 3,000 x *g* (15 min, 25°C, with no brake) and sequentially passed through filters with 5 μm, 1.2 μm, and 0.45 μm cutoffs. The pellets containing protoplast cells were stored at −20°C for sterol quantification. After filtration, the supernatants were concentrated in an Amicon ultrafiltration system (cutoff, 100 kDa) and again centrifuged at 10,000 x *g*, 4°C for 15 min to eliminate possible cellular debris. The concentrated supernatants were finally ultracentrifuged at 100,000 x *g* (4°C, 1 h). The resulting pellets containing EVs were washed twice with 0.22 μm-filtered PBS under the same ultracentrifugation conditions and finally suspended in 300 μl of 0.22 μm-filtered PBS. EVs suspensions were stored at −80°C for further experiments.

For nanoparticle tracking analysis (NTA) and GAG serological detection, the EV suspensions were prior submitted to immunoprecipitation for the removal of non-vesicular polysaccharides. In this assay, 50 μl of the EVs suspension was added to the wells of a 96-well enzyme-linked immunosorbent assay (ELISA) plate, previously coated with a mixture of antibodies against α-glucan (J558) (64), β-glucan (Dectin1-human IgG Fc chimeric β-(1,3)-glucan receptor; a kind gift of G. Brown, University of Aberdeen, United Kingdom) and GAG (1 μg/ml, 1 h, 37°C) (44) and blocked with PBS containing 1 % bovine serum albumin (BSA). Unbound fractions were collected and the resulting EVs were stored at −80°C for further experiments.

Nanoparticle tracking analysis (NTA) was performed to determine the EV diameter and concentration. NTA of protoplast EVs was performed on an LM10 nanoparticle analysis system, coupled with a 488-nm laser and equipped with an SCMOS camera and a syringe pump (Malvern Panalytical, Malvern, United Kingdom), as recently described for *C. gattii* EVs (65). The samples were 25-fold diluted in filtered PBS and measured within the optimal dilution range of 7,6 ×10^7^ to 6.8 ×10^8^ particles/ml. The data were acquired and analyzed using the NTA 3.0 software (Malvern Panalytical). The quantification of sterol in EV preparations was performed with the Amplex^®^ Red Sterol Assay Kit (14, 66, 67).

### Transmission electron microscopy (TEM) of EVs

For negative staining TEM, the EV pellets were fixed with 2% glutaraldehyde, 2% paraformaldehyde in 0.1 M sodium cacodylate buffer at room temperature for 2 h and then post-fixed overnight at 4°C with 1% glutaraldehyde, 4% paraformaldehyde in PBS. Copper carbon coated grids (Cu-CF300, EMS), previously negatively charged with the ELMO system (1 min, 0,15 mbar 2mA 80 volts) were put in contact with 15 μl of each sample for 10 min, and washed three times with Milli-Q water drops (2 min each), stained with uranyl acetate 2%, dried and observed with a Tecnai Spirit microscope operating at 120 kV and equipped with an EAGLE 4kx4k camera.

EVs were alternatively fixed with 2% formaldehyde with 2% glutaraldehyde in cacodylate buffer (0.1M, pH 7.4). The samples were washed through four changes of cacodylate buffer (30 min each) and pelleted in 1% agarose (JT Baker Chemical Co., Phillipsburg, New Jersey). They were transferred to 1% osmium tetroxide in cacodylate buffer (0.1 M, pH 7.4) and incubated at 4 °C for 1 h followed by washing in cacodylate buffer and distilled water for a total of 30 min. The samples were then stained with 0.5% aqueous uranyl acetate, dehydrated, slowly infiltrated with epoxy and embedded. After resin polymerization, the blocks were sectioned on a Leica ultramicrotome and poststained for 10 min in 2% uranyl acetate in 50% ethanol and for 5 min in lead citrate. Ultrathin sections (70 nm thick) were collected on formvar-coated copper slot grids and poststained for 10 min in 2% uranyl acetate in 50% ethanol and for 5 min in lead citrate. Sections were then examined on a JEOL 1200EX (JEOL Ltd., Tokyo, Japan) microscope equipped with a SIA L3C CCD camera (SIA Inc., Duluth, Georgia).

### Monosaccharide composition and serological detection of GAG in EVs

EV ultracentrifugation pellets were suspended in water for monosaccharide analysis. Monosaccharides present in the EVs were determined by gas chromatography after hydrolysis, reduction and paracetylation of the vesicle components using meso-inositol as internal standard (69). Serological estimation of vesicular GAG was performed as described before by our group for other fungal polysaccharides (67). Briefly, EV suspensions were vacuum dried and suspended in chloroform:methanol (9:1, vol/vol). The suspension was centrifuged, and the resulting white precipitate was solubilized in PBS for quantitative ELISA with the anti-GAG antibody. The purified polysaccharide (SGG, a kind gift of T. Fontaine, Institut Pasteur, Paris, France) was used for the preparation of standard curves and determination of polysaccharide concentrations in EV samples (44).

### Protein composition of EVs

EV ultracentrifugation pellets were suspended in buffer containing 8 M urea in 100 mM Tris, pH 7.5, for proteomic analysis. Protein samples were reduced with 5 mM dithiothreitol (DTT) for 30 min at 23°C and then alkylated with 20 mM iodoacetamide in the dark at room temperature, for 30 min. Subsequently, the endoproteinase LysC (Promega) was added for the first digestion step (protein to Lys-C ratio = 80:1) for 3 h at 30°C. Then the sample was diluted to reach the 1 M urea concentration with 100 mM Tris (pH7.5), and trypsin (Promega) was added to the sample (protein to trypsin ratio = 50:1). The samples were digested for 16 h at 37°C. Proteolysis was stopped by the addition of 1% formic acid. The resulting peptides were desalted using a Sep-Pak SPE cartridge (Waters) according to manufacturer’s instructions.

Liquid chromatography coupled to mass spectrometry (LC-MS/MS) analysis of digested peptides was performed on an Orbitrap Q Exactive Plus mass spectrometer (Thermo Fisher Scientific, Bremen) coupled to an EASY-nLC 1000 (Thermo Fisher Scientific). The peptides were loaded and separated at 250 nl.min-1 on a home-made C18 50 cm capillary column with a picotip silica emitter (75 μm diameter filled with 1.9 μm Reprosil-Pur Basic C18-HD resin, Dr. Maisch GmbH, Ammerbuch-Entringen, Germany) equilibrated in solvent A (0.1 % formic acid). The peptides were eluted using a gradient of solvent B (Acetonitrile (ACN), 0.1 % FA) from 2 to 18 % for 110 min, 18 to 30 % for 35 min, 30 to 45 % for 15 min under a 250 nl/min flow rate. The total length of the chromatographic run was 185 min, including high ACN level steps and column regeneration. Mass spectra were obtained in the data-dependent acquisition mode with the Xcalibur 2.2 software (Thermo Fisher Scientific, Bremen) with automatic switching between MS and MS/MS scans using a top-10 method. Spectral resolution corresponded to 70,000 (at m/z 400) with a target value of 3 × 10^6^ ions. The scan range was limited from 300 to 1,700 m/z. Peptide fragmentation was performed via higher-energy collision dissociation (HCD) with the energy set at 28 normalized collision energy (NCE). Intensity threshold for the ion selection was set at 1 × 10^6^ ions with charge exclusion of z = 1 and z > 7. The MS/MS spectra were acquired at a resolution of 17,500 (at m/z 400). The isolation window was set at 1.6 Th. Dynamic exclusion was employed within 45s.

Data search was performed with MaxQuant tool (70) (version 1.5.3.8) with the Andromeda search engine against the *A. fumigatus* A1163 database (9942 entries, downloaded from https://www.uniprot.org in September 18, 2019. As search parameters, carbamidomethylation of cysteines was set as a fixed modification, oxidation of methionine and protein N-terminal acetylation were set as variable modifications. The mass tolerances in MS and MS/MS were set to 5 ppm and 20 ppm, respectively. Maximum peptide charge was set to 7 and 7 amino acids were required as the minimum peptide length. A false discovery rate of 1% was set for both protein and peptide levels. Four independent EV ultracentrifugation pellets were analyzed for protoplasts submitted either to cell wall regenerating and repressing conditions. Two EV samples of freshly purified protoplasts were analyzed. Proteins identified in at least two independent experiments were assigned for Gene Ontology (GO). GO term enrichment was performed using the GOtermfinder feature of the AspGD database.(http://www.aspergillusgenome.org/cgi-bin/GO/goTermFinder). Enriched GO terms were summarized by removing redundancy using the REVIGO web server, available at http://revigo.irb.hr/ (71). Revigo output were viewed in R (v4.0.0) using the treemap package.

### Statistical analysis

All statistical analyses were performed using the GraphPad Prism 6 software (GraphPad Software Inc.). Data sets were tested for normal distribution using a Shapiro–Wilk or Kolmogorov-Smirnov normality tests. When passed the normality test (alpha=0.05), the data were further analyzed using the unpaired Student’s t-test. Multiple data sets were further analyzed using ordinary one-way ANOVA, followed by the Tukey’s multiple comparison test. When at least one data set was non-normally distributed, multiple data sets were analyzed by the nonparametric Kruskal–Wallis’ test.

## References

1. Krappmann S. 2016. How to invade a susceptible host: cellular aspects of aspergillosis. Curr Opin Microbiol.

2. Abad A, Victoria Fernández-Molina J, Bikandi J, Ramírez A, Margareto J, Sendino J, Luis Hernando F, Pontón J, Garaizar J, Rementeria A. 2010. What makes Aspergillus fumigatus a successful pathogen? Genes and molecules involved in invasive aspergillosis. Rev Iberoam Micol.

3. Sugui JA, Kwon-Chung KJ, Juvvadi PR, Latgé JP, Steinbach WJ. 2015. Aspergillus fumigatus and related species. Cold Spring Harb Perspect Med.

4. Latgé JP, Chamilos G. 2020. Aspergillus fumigatus and aspergillosis in 2019. Clin Microbiol Rev.

5. Casadevall A, Nosanchuk JD, Williamson P, Rodrigues ML. 2009. Vesicular transport across the fungal cell wall. Trends Microbiol 17.

6. Wolf JM, Casadevall A. 2014. Challenges posed by extracellular vesicles from eukaryotic microbes. Curr Opin Microbiol.

7. Brown L, Wolf JM, Prados-Rosales R, Casadevall A. 2015. Through the wall: extracellular vesicles in Gram-positive bacteria, mycobacteria and fungi. Nat Rev Microbiol 13:620.

8. Nimrichter L, De Souza MM, Del Poeta M, Nosanchuk JD, Joffe L, Tavares PDM, Rodrigues ML. 2016. Extracellular vesicle-associated transitory cell wall components and their impact on the interaction of fungi with host cells. Front Microbiol.

9. Rodrigues ML, Nimrichter L, Oliveira DL, Frases S, Miranda K, Zaragoza O, Alvarez M, Nakouzi A, Feldmesser M, Casadevall A. 2007. Vesicular polysaccharide export in Cryptococcus neoformans is a eukaryotic solution to the problem of fungal trans-cell wall transport. Eukaryot Cell 6:48–59.

10. Bielska E, Sisquella MA, Aldeieg M, Birch C, O’Donoghue EJ, May RC. 2018. Pathogen-derived extracellular vesicles mediate virulence in the fatal human pathogen Cryptococcus gattii. Nat Commun 9.

11. Albuquerque PC, Nakayasu ES, Rodrigues ML, Frases S, Casadevall A, Zancope-Oliveira RM, Almeida IC, Nosanchuk JD. 2008. Vesicular transport in Histoplasma capsulatum: An effective mechanism for trans-cell wall transfer of proteins and lipids in ascomycetes. Cell Microbiol 10.

12. Gehrmann U, Qazi KR, Johansson C, Hultenby K, Karlsson M, Lundeberg L, Gabrielsson S, Scheynius A. 2011. Nanovesicles from malassezia sympodialis and host exosomes induce cytokine responses - novel mechanisms for host-microbe interactions in atopic eczema. PLoS One.

13. Vallejo MC, Matsuo AL, Ganiko L, Medeiros LCS, Miranda K, Silva LS, Freymüller-Haapalainen E, Sinigaglia-Coimbra R, Almeida IC, Puccia R. 2011. The pathogenic fungus Paracoccidioides brasiliensis exports extracellular vesicles containing highly Immunogenic α-galactosyl epitopes. Eukaryot Cell.

14. Vargas G, Rocha JDB, Oliveira DL, Albuquerque PC, Frases S, Santos SS, Nosanchuk JD, Gomes AMO, Medeiros LCAS, Miranda K, Sobreira TJP, Nakayasu ES, Arigi EA, Casadevall A, Guimaraes AJ, Rodrigues ML, Freire-de-Lima CG, Almeida IC, Nimrichter L. 2015. Compositional and immunobiological analyses of extracellular vesicles released by Candida albicans. Cell Microbiol 17.

15. Leone F, Bellani L, Muccifora S, Giorgetti L, Bongioanni P, Simili M, Maserti B, Del Carratore R. 2017. Analysis of extracellular vesicles produced in the biofilm by the dimorphic yeast Pichia fermentans. J Cell Physiol 233:2759–2767.

16. Augusto M, Ikeda K, Roberto J, Almeida F De, Jannuzzi GP, Cronemberger-andrade A, Pinheiro J, Almeida SR De, Miranda DZ. 2018. Extracellular Vesicles From Sporothrix brasiliensis Are an Important Virulence Factor That Induce an Increase in Fungal Burden in Experimental Sporotrichosis. Front Microbiol.

17. Lavrin T, Konte T, Kostanjsek R, Sitar S, Sepcic K, Prpar Mihevc S, Zagar E, Zupunski V, Lenassi M, Rogelj B, Gunde Cimerman N. 2020. The Neurotropic Black Yeast Exophiala dermatitidis Induces Neurocytotoxicity in Neuroblastoma Cells and Progressive Cell Death. Cells 9:963.

18. Silva BMA, Prados-Rosales R, Espadas-Moreno J, Wolf JM, Luque-Garcia JL, Gonçalves T, Casadevall A. 2014. Characterization of Alternaria infectoria extracellular vesicles. Med Mycol.

19. Bleackley MR, Samuel M, Garcia-Ceron D, McKenna JA, Lowe RGT, Pathan M, Zhao K, Ang CS, Mathivanan S, Anderson MA. 2020. Extracellular Vesicles From the Cotton Pathogen Fusarium oxysporum f. sp. vasinfectum Induce a Phytotoxic Response in Plants. Front Plant Sci.

20. Bitencourt TA, Rezende CP, Quaresemin NR, Moreno P, Hatanaka O, Rossi A, Martinez-Rossi NM, Almeida F. 2018. Extracellular vesicles from the dermatophyte trichophyton interdigitalemodulate macrophage and keratinocyte functions. Front Immunol.

21. Liu M, Bruni GO, Taylor CM, Zhang Z, Wang P. 2018. Comparative genome-wide analysis of extracellular small RNAs from the mucormycosis pathogen Rhizopus delemar. Sci Rep.

22. Souza JAM, Baltazar L de M, Carregal VM, Gouveia-Eufrasio L, de Oliveira AG, Dias WG, Rocha de Miranda K, Malavazi I, Santos D de A, Frézard FJG, de Souza D da G, Teixeira MM, Soriani FM. 2019. Characterization of Aspergillus fumigatus Extracellular Vesicles and Their Effects on Macrophages and Neutrophils Functions. Front Microbiol.

23. Gibson RK, Peberdy JF. 1972. Fine structure of protoplasts of Aspergillus nidulans. J Gen Microbiol.

24. Raposo G, Stoorvogel W. 2013. Extracellular vesicles: Exosomes, microvesicles, and friends. J Cell Biol.

25. Peberdy JF, Gibson RK. 1971. Regeneration of Aspergillus nidulans protoplasts. J Gen Microbiol.

26. Osumi M. 1998. The ultrastructure of yeast: Cell wall structure and formation. Micron.

27. Pardo M, Monteoliva L, Pla J, Sánchez M, Gil C, Nombela C. 1999. Two-dimensional analysis of proteins secreted by Saccharomyces cerevisiae regenerating protoplasts: A novel approach to study the cell wall. Yeast.

28. Pitarch A, Nombela C, Gil C. 2008. Collection of proteins secreted from yeast protoplasts in active cell wall regeneration. Methods Mol Biol.

29. Gil-Bona A, Reales-Calderon JA, Parra-Giraldo CM, Martinez-Lopez R, Monteoliva L, Gil C. 2016. The cell wall protein Ecm33 of Candida albicans is involved in chronological life span, morphogenesis, cell wall regeneration, stress tolerance, and host-cell interaction. Front Microbiol.

30. Rodrigues ML, Nakayasu ES, Oliveira DL, Nimrichter L, Nosanchuk JD, Almeida IC, Casadevall A. 2008. Extracellular vesicles produced by Cryptococcus neoformans contain protein components associated with virulence. Eukaryot Cell 7.

31. Beauvais A, Bruneau JM, Mol PC, Buitrago MJ, Legrand R, Latgé JP. 2001. Glucan synthase complex of Aspergillus fumigatus. J Bacteriol.

32. Henry C, Latgé JP, Beauvais A. 2012. α1,3 glucans are dispensable in Aspergillus fumigatus. Eukaryot Cell.

33. Muszkieta L, Aimanianda V, Mellado E, Gribaldo S, Alcàzar-Fuoli L, Szewczyk E, Prevost MC, Latgé JP. 2014. Deciphering the role of the chitin synthase families 1 and 2 in the in vivo and in vitro growth of Aspergillus fumigatus by multiple gene targeting deletion. Cell Microbiol.

34. Henry C, Li J, Danion F, Alcazar-Fuoli L, Mellado E, Beau R, RémiJouvion G, Latgé JP, Fontainea T. 2019. Two KTR mannosyltransferases are responsible for the biosynthesis of cell wall mannans and control polarized growth in aspergillus fumigatus. MBio.

35. Briard B, Muszkieta L, Latgé JP, Fontaine T. 2016. Galactosaminogalactan of Aspergillus fumigatus, a bioactive fungal polymer. Mycologia.

36. Mouyna I, Hartl L, Latgé JP. 2013. β-1,3-glucan modifying enzymes in Aspergillus fumigatus. Front Microbiol.

37. Henry C, Fontaine T, Heddergott C, Robinet P, Aimanianda V, Beau R, Beauvais A, Mouyna I, Prevost MC, Fekkar A, Zhao Y, Perlin D, Latgé JP. 2016. Biosynthesis of cell wall mannan in the conidium and the mycelium of Aspergillus fumigatus. Cell Microbiol.

38. Mouyna I, Kniemeyer O, Jank T, Loussert C, Mellado E, Aimanianda V, Beauvais A, Wartenberg D, Sarfati J, Bayry J, Prévost MC, Brakhage AA, Strahl S, Huerre M, Latgé JP. 2010. Members of protein O-mannosyltransferase family in Aspergillus fumigatus differentially affect growth, morphogenesis and viability. Mol Microbiol.

39. Rodrigues ML, Oliveira DL, Vargas G, Girard-Dias W, Franzen AJ, Frasés S, Miranda K, Nimrichter L. 2016. Analysis of yeast extracellular vesiclesMethods in Molecular Biology.

40. Selitrennikoff CP, Bloomfield EC. 1984. Formation and regeneration of protoplasts of wild-type Neurospora crassa. Curr Microbiol.

41. Wolf JM, Espadas-Moreno J, Luque-Garcia JL, Casadevall A. 2014. Interaction of cryptococcus neoformans extracellular vesicles with the Cell Wall. Eukaryot Cell.

42. Hochstenbach F, Klis FM, Van Ende H Den, Van Donselaar E, Peters PJ, Klausner RD. 1998. Identification of a putative alpha-glucan synthase essential for cell wall construction and morphogenesis in fission yeast. Proc Natl Acad Sci U S A.

43. Sánchez-León E, Riquelme M. 2015. Live imaging of β-1,3-glucan synthase FKS-1 in Neurospora crassa hyphae. Fungal Genet Biol.

44. Fontaine T, Delangle A, Simenel C, Coddeville B, van Vliet SJ, van Kooyk Y, Bozza S, Moretti S, Schwarz F, Trichot C, Aebi M, Delepierre M, Elbim C, Romani L, Latgé JP. 2011. Galactosaminogalactan, a new immunosuppressive polysaccharide of Aspergillus fumigatus. PLoS Pathog.

45. Gresnigt MS, Bozza S, Becker KL, Joosten LAB, Abdollahi-Roodsaz S, van der Berg WB, Dinarello CA, Netea MG, Fontaine T, De Luca A, Moretti S, Romani L, Latge JP, van de Veerdonk FL. 2014. A Polysaccharide Virulence Factor from Aspergillus fumigatus Elicits Anti-inflammatory Effects through Induction of Interleukin-1 Receptor Antagonist. PLoS Pathog.

46. Loussert C, Schmitt C, Prevost MC, Balloy V, Fadel E, Philippe B, Kauffmann-Lacroix C, Latgé JP, Beauvais A. 2010. In vivo biofilm composition of Aspergillus fumigatus. Cell Microbiol.

47. Rodrigues ML, Nimrichter L, Oliveira DL, Frases S, Miranda K, Zaragoza O, Alvarez M, Nakouzi A, Feldmesser M, Casadevall A. 2007. Vesicular polysaccharide export in Cryptococcus neoformans is a eukaryotic solution to the problem of fungal trans-cell wall transport. Eukaryot Cell 6.

48. Schuster M, Martin-Urdiroz M, Higuchi Y, Hacker C, Kilaru S, Gurr SJ, Steinberg G. 2016. Co-Delivery of Cell-Wall-Forming enzymes in the same vesicle for coordinated fungal cell wall formation. Nat Microbiol.

49. Beauvais A, Perlin DS, Latgé JP. 2007. Role of α(1-3)-glucan in Aspergillus fumigatus and other human fungal pathogens, p. 269–288. In Gadd, G, Dyer, PS, Watkinson, SC (eds.), Fungi in the Environment. Cambridge University Press, Cambridge.

50. Gastebois A, Fontaine T, Latgé JP, Mouyna I. 2010. β(1-3)glucanosyltransferase Gel4p is essential for Aspergillus fumigatus. Eukaryot Cell.

51. Aimanianda V, Simenel C, Garnaud C, Clavaud C, Tada R, Barbin L, Mouyna I, Heddergott C, Popolo L, Ohya Y, Delepierre M, Latgea JP. 2017. The dual activity responsible for the elongation and branching of β-(1,3)-glucan in the fungal cell wall. MBio.

52. Dichtl K, Samantaray S, Aimanianda V, Zhu Z, Prévost MC, Latgé JP, Ebel F, Wagener J. 2015. Aspergillus fumigatus devoid of cell wall β-1,3-glucan is viable, massively sheds galactomannan and is killed by septum formation inhibitors. Mol Microbiol.

53. Rodrigues ML, Franzen AJ, Nimrichter L, Miranda K. 2013. Vesicular mechanisms of traffic of fungal molecules to the extracellular space. Curr Opin Microbiol 16.

54. Asif AR, Oellerich M, Amstrong VW, Riemenschneider B, Monod M, Reichard U. 2006. Proteome of conidial surface associated proteins of Aspergillus fumigatus reflecting potential vaccine candidates and allergens. J Proteome Res.

55. Champer J, Diaz-Arevalo D, Champer M, Hong TB, Wong M, Shannahoff M, Ito JI, Clemons K V., Stevens DA, Kalkum M. 2012. Protein targets for broad-spectrum mycosis vaccines: Quantitative proteomic analysis of Aspergillus and Coccidioides and comparisons with other fungal pathogens. Ann N Y Acad Sci.

56. Karkowska-Kuleta J, Kozik A. 2015. Cell wall proteome of pathogenic fungi. Acta Biochim Pol.

57. Champer J, Ito JI, Clemons K V., Stevens DA, Kalkum M. 2016. Proteomic analysis of pathogenic fungi reveals highly expressed conserved cell wall proteins. J Fungi.

58. Da Silva Ferreira ME, Kress MRVZ, Savoldi M, Goldman MHS, Härtl A, Heinekamp T, Brakhage AA, Goldman GH. 2006. The akuBKU80 mutant deficient for nonhomologous end joining is a powerful tool for analyzing pathogenicity in Aspergillus fumigatus. Eukaryot Cell.

59. Cove DJ. 1966. The induction and repression of nitrate reductase in the fungus Aspergillus nidulans. Biochim Biophys Acta.

60. Briard B, Bomme P, Lechner BE, Mislin GLA, Lair V, Prévost MC, Latgé JP, Haas H, Beauvais A. 2015. Pseudomonas aeruginosa manipulates redox and iron homeostasis of its microbiota partner Aspergillus fumigatus via phenazines. Sci Rep.

61. Beauvais A, Bozza S, Kniemeyer O, Formosa C, Balloy V, Henry C, Roberson RW, Dague E, Chignard M, Brakhage AA, Romani L, Latgé JP. 2013. Deletion of the α-(1,3)-Glucan Synthase Genes Induces a Restructuring of the Conidial Cell Wall Responsible for the Avirulence of Aspergillus fumigatus. PLoS Pathog.

62. Ramos CL, Gomes FM, Girard-Dias W, Almeida FP, Albuquerque PC, Kretschmer M, Kronstad JW, Frases S, De Souza W, Rodrigues ML, Miranda K. 2017. Phosphorus-rich structures and capsular architecture in Cryptococcus neoformans. Future Microbiol 12:227–238.

63. Paris S, Debeaupuis JP, Crameri R, Carey M, Charlès F, Prévost MC, Schmitt C, Philippe B, Latgé JP. 2003. Conidial hydrophobins of Aspergillus fumigatus. Appl Environ Microbiol.

64. Kearney JF, McCarthy MTM, Stohrer R, Benjamin WH, Briles DE. 1985. Induction of germ-line anti-α1-3 dextran antibody responses in mice by members of the enterobacteriaceae family. J Immunol.

65. Reis FCG, Borges BS, Jozefowicz LJ, Sena BAG, Garcia AWA, Medeiros LC, Martins ST, Honorato L, Schrank A, Vainstein MH, Kmetzsch L, Nimrichter L, Alves LR, Staats CC, Rodrigues ML. 2019. A Novel Protocol for the Isolation of Fungal Extracellular Vesicles Reveals the Participation of a Putative Scramblase in Polysaccharide Export and Capsule Construction in Cryptococcus gattii mSphere 4:e00080–19.

66. Oliveira DL, Nakayasu ES, Joffe LS, Guimarães AJ, Sobreira TJP, Nosanchuk JD, Cordero RJB, Frases S, Casadevall A, Almeida IC, Nimrichter L, Rodrigues ML. 2010. Characterization of yeast extracellular vesicles: Evidence for the participation of different pathways of cellular traffic in vesicle biogenesis. PLoS One 5.

67. Rizzo J, Oliveira DL, Joffe LS, Hu G, Gazos-Lopes F, Fonseca FL, Almeida IC, Frases S, Kronstad JW, Rodrigues ML. 2014. Role of the Apt1 protein in polysaccharide secretion by Cryptococcus neoformans. Eukaryot Cell 13:715–726.

68. Folch J, Lees M, Sloane Stanley GH. 1957. A simple method for the isolation and purification of total lipides from animal tissues. J Biol Chem.

69. Sawardeker JS, Sloneker JH, Jeanes A. 1965. Quantitative determination of monosaccharides and their acetates by gas liquid chromatography. Anal Chem F Full J Title.

70. Tyanova S, Temu T, Cox J. 2016. The MaxQuant computational platform for mass spectrometry-based shotgun proteomics. Nat Protoc.

71. Supek F, Bošnjak M, Škunca N, Šmuc T. 2011. Revigo summarizes and visualizes long lists of gene ontology terms. PLoS One.

